# Pyroptosis of syncytia formed by fusion of SARS-CoV-2 Spike and ACE2 expressing cells

**DOI:** 10.1101/2021.02.25.432853

**Authors:** Huabin Ma, Zhoujie Zhu, Huaipeng Lin, Shanshan Wang, Peipei Zhang, Yanguo Li, Long Li, Jinling Wang, Yufen Zhao, Jiahuai Han

**Affiliations:** Institute of Drug Discovery Technology, Ningbo University, Ningbo, Zhejiang 315211, China; Qian Xuesen Collaborative Research Center of Astrochemistry and Space Life Sciences, Ningbo University, Ningbo, Zhejiang 315211, China; State Key Laboratory of Cellular Stress Biology, School of Life Sciences, Xiamen University, Xiamen, Fujian 361102, China; Research Unit of Cellular Stress of CAMS, Cancer Research Center of Xiamen University, Xiang’an Hospital of Xiamen University, School of Medicine, Xiamen University, Xiamen, Fujian 361102, China; Department of Emergency, Zhongshan Hospital of Xiamen University, Xiamen 361005, China

**Author notes:** **Correspondence**: Jiahan Han;, Yufen Zhao;, Huabin Ma.

## Abstract

SARS-Cov-2 infected cells fused with the ACE2-positive neighboring cells forming syncytia. However, the effect of syncytia in disease development is largely unknown. We established an *in vitro* cell-cell fusion system and used it to mimic the fusion of SARS-CoV-2 infected cells with ACE2-expressing cells to form syncytia. We found that Caspase-9 was activated after syncytia formation, and Caspase-3/7 was activated downstream of Caspase-9, but it triggered GSDME-dependent pyroptosis rather than apoptosis. What is more, single cell RNA-sequencing data showed that both ACE2 and GSDME were expression in alveolar type 2 cells in human lung. We propose that pyroptosis is the fate of syncytia formed by SARS-CoV-2 infected host cells and ACE2-positive cells, which indicated that lytic death of syncytia may contribute to the excessive inflammatory responses in severe COVID-19 patients.

## Results

CoronaVirus Disease 2019 (COVID-19) is an infectious disease associated with systematical multi-organ failure caused by SARS-CoV-2, which mainly infects the lung and upper respiratory system^1,2^. Massive multinucleated syncytia are commonly observed in autopsy of severe COVID-19 patients^3^. It has been reported that the interaction between Spike (S) protein and ACE2 not only mediated the fusion of virus with host cells, but also multinucleated giant cells formation^4,5^. However, the effect of syncytia in disease development is largely unknown.

In order to better observe the formation of syncytia, we established an *in vitro* cell-cell fusion system and used it to mimic the fusion of SARS-CoV-2 infected cells with ACE2-expressing cells to form syncytia. High-content imaging with confocal laser scanning microscope was used to characterize the SARS-CoV-2-S-GFP expressing cells, which showed that SARS-CoV-2-S-GFP was mainly on the cell surface and was co-localized with cell membrane marker PLCδ-PH (Fig. S1a) but not nuclear RFP (Fig. S1e)^6^. In addition, S protein was detected as puncta in cytoplasm with co-localization of Golgi marker GGA1 (Fig. S1b), consisting with reports showing that the S protein was glycosylated in Golgi apparatus^7^. Two endosome-related proteins Rab5a and Rab7a were also co-located with the S protein, indicating the trafficking of S protein was associated with cellular membrane system (Fig. S1c and d). Then, the SARS-CoV-2-S-GFP expressing cell lines, named 2S-GFP-HeLa, were co-cultured with A549 expressing ACE2 cells (PLC-RFP-A549+ACE2) at a 1:1 ratio. Syncytia were observed 4 hours later (Fig. 1a), and the cell-cell fusion occurred between the cell membranes because of the nuclei were intact (Fig. 1b). Additionally, syncytia were formed in co-culture system regardless of whether they were the same type of cells (2S-GFP-A549 and A549+ACE2, Fig. S2a) or different types of cells (2S-GFP-H1299 and A549+ACE2, Fig. S2b; 2S-GFP-HeLa and A549+ACE2, Fig.1a), or even cells from different species (2S-GFP-L929 and A549+ACE2, Fig. S2c).

**Fig.1.**
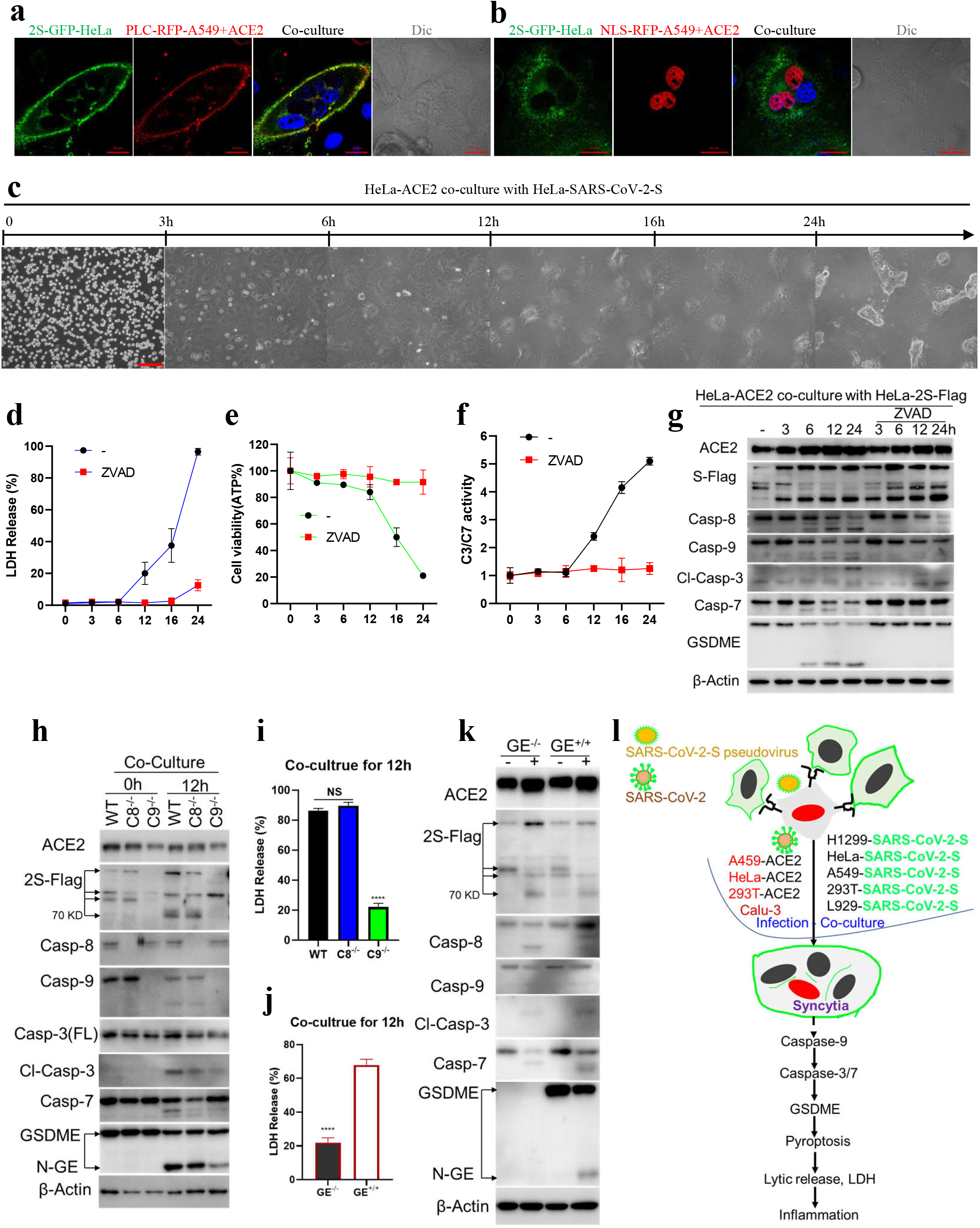
Caspase-9/GSDME trigged pyroptosis of syncytia formed by fusion of SARS-CoV-2 Spike and ACE2 expressing cells. SARS-CoV-2-S-GFP HeLa cells were co-cultured with PLC-RFP-A549+ACE2 (**a**), or NLS-RFP-A549+ACE2 cells (**b**) at a 1:1 ratio, four hours later, image was scanned by LSM780, the nucleus(blue) was stained by Hoechst; Bar, 20 μm. **c**. The time-phase of cell-cell fusion progress, HeLa-ACE2 co-culture with HeLa-SARS-CoV-2-S cells, images were obtained by Digital Cell Checker for Cell Culture PAULA; Bar, 200 μm; then the LDH release (**d**),ATP (**e**),the activity of Caspase-3/7 (**f**) were detected at indicated time,pan-caspase inhibitor, ZVAD(20 uM) was used. LDH release was measured after co-culture for 12 hours as indicated (**i** and **j**). Western blot analysis of the cells collected as indicated, ACE2, Flag, Caspase-8/9/3/7, and GSMDE were probed, β-Actin as loading control (**g, h** and **k**). The model of SARS-CoV-2 Spike interaction with ACE2 induced syncytia pyroptosis (**l**). The data were shown as means ± SD,****P<0.0001, NS, non-significance, and the experiments were replicated more than three times.

It is well known that SARS-CoV-2 and SARS-CoV-1 enter host cells through the same receptor ACE2^8^. To find out whether SARS-CoV-1-S can interact with ACE2 and lead to cell-cell fusion like SARS-CoV-2-S. 293T-ACE2 cells were co-cultured with HeLa cells expressing SARS-CoV-1-S-Flag, or SARS-CoV-2-S-Flag. SARS-CoV-1-S-Flag HeLa cells indeed fused with 293T-ACE2 cells, while the degree of fusion is lower compared with SARS-CoV-2-S-Flag cells (Fig. S2d), even if the expression of S protein of SARS-CoV-1 is higher than SARS-CoV-2 (Fig. S2e). The similar results were obtained when HeLa cells expressing SARS-CoV-1-S-Flag, or SARS-CoV-2-S-Flag were co-cultured with Calu-3, which is a human lung cancer epithelioid cell line expressing endogenous ACE2 and can be infected by SARS-CoV-2 (Fig. S2f and g). Thus, the degree of cell-cell fusion mediated by S protein of SARS-CoV-2 interaction with ACE2 was higher than SARS-CoV-1, which might due to difference in the affinity to ACE2 between the S proteins of SARS-CoV-1 and SARS-CoV-2.

Next, we aimed to investigate the fate of syncytia. To eliminate the cross-talk of different cytosolic contents between different cell types, we co-cultured HeLa cells expressing SARS-CoV-2-S (HeLa-2S) with HeLa-ACE2, and recorded the process of cell fusion by the real-time observation system (Fig. 1c and Movie-1). We found that syncytia formed upon cell-cell fusion, grew in size steadily, finally, ruptured with LDH release, ATP decrease, and the activity of caspase-3/7 increase. The death was blocked by pan-caspase inhibitor zVAD (Fig. 1d, e and f). We assessed the activation of molecules related to the death. As shown in Fig. 1g, caspase-8/9 and caspase-3/7 were cleaved, suggesting an activation of apoptosis pathway. Interestingly, we also detected cleavage of GSDME. It is known that activation of GSDME is mediated by caspase-3^9,10^, and we confirmed the role of caspase in activation of GSDME as zVAD effectively blocked GSDME cleavage. Thus, it can be proposed that syncytia formation led to activation of caspase-8/9 to caspase3/7 cascade. The activated caspase-3 cleaved GSDME and released N-GSDME to permeabilize cell membrane to execute syncytia pyroptosis.

To define the death pathway underlying syncytia pyroptosis, we knocked out caspase-8 (*Casp8*^*-/-*^) or caspase-9 (*Casp9*^*-/-*^) in HeLa cells by CRISPR-Cas9 and expressed ACE2 and SARS-CoV-2-S-Flag in these cells respectively, and then co-cultured these cells in pairs. We observed syncytia formation in *Casp8*^*-/-*^ and *Casp9*^*-/-*^ cells without difference from WT cells (Fig. S3a). In contrast, *Casp9* but not *Casp8* deletion blocked syncytia death (Fig. 1i). Further analysis showed that the cleavage of GSDME (N-GE) was significantly decreased in *Casp9*^*-/-*^ cells (Fig. 1h). Unexpectedly, the 70kD fragment of SARS-CoV-2-S-Flag, which was believed to be processed by protease TMPRSS2 or Cathepsin L during S protein fusion with ACE2, was disappeared in *Casp9*^*-/-*^ cells, indicating a linkage between caspase-9 and SARS-CoV-2 S protein cleavage.

To further confirm that GSDME was involved in the death of syncytia, we generated GSDME knock-out (*GE*^*-/-*^+ACE2, *GE*^*-/-*^+SARS-CoV-2-S-Flag) and WT HeLa cell lines (*GE*^*+/+*^+ACE2, *GE*^*+/+*^+SARS-CoV-2-S-Flag), and co-cultured them. Similarly, GSDME did not affect cell fusion to form syncytia (Fig. S3b and c), but GSDME knock-out significantly inhibited the death of syncytia (Fig. 1j). In addition, GSDME knock-out did not affect the activation of caspases (Fig. 1k), confirming it is downstream of caspases.

Our data indicated that the death of syncytia induced by SARS-CoV-2 infection could be mediated by GSDME-dependent pyroptosis. The existing transcriptomic data^11^ showed correlations of the expression between ACE2 and GSDME, especially in the small intestine and testis (Fig. S4). The high-level expression of ACE2 and GSDME in testis could link to the destruction of male reproductive system by SARS-CoV-2 infection^12^. We analyzed single-cell-RNA-sequencing (scRNA-Seq) data from eight normal human lung transplant donors with a total of 42,225 cells^13^. As reported, the expression of ACE2 is concentrated in a special small population of alveolar type 2 (AT2) cells in lung^14^ (Fig. S5a), and GSDME is also enrichment in AT2 cells. Thus, GSDME-dependent pyroptosis could occur in SARS-CoV-2 infected AT2 cells (Fig. S5b).

Here, we provide *in vitro* evidence showing that the syncytia formed by fusion of the cells expressing SARS-CoV-2 S protein and ACE2 respectively undergo pyroptosis (Fig. 1l). The pyroptosis is initiated by components of intrinsic apoptosis pathway and executed by caspase-3/7 mediated activation/cleavage of GSDME. Since scRNA-seq data showed that both ACE2 and GSDME were expression in AT2 cells in human lung, we propose that pyroptosis is the fate of syncytia formed by SARS-CoV-2 infected host cells and ACE2-positive cells. The lytic death of syncytia may contribute to the excessive inflammatory responses in severe COVID-19 patients.

## ACKNOWLEDGEMENTS

We thank Peihui Wang (Shandong university) for kindly providing the plasmid of SARS-CoV-2-S. We thank Lu Zhou (Xiamen university) and Yumei Xie (Fujian Jiangxia university) for scientific editing of the manuscript.

This work was supported by grants from the National Natural Science Foundation of China (81788101, 81630042 to J.H.; 81700596 to H.M.; 91856126 to Y.Z.).

## AUTHOR CONTRIBUTIONS

J.H., Y.Z., H.M. conceived and designed the experiments. H.M., Z.Z., H.L., P.Z., L.L. performed the experiments: generated the cell strains and performed immunofluorescent imaging, and the associated western blot analyses. S.W. and Y.L. analyzed the data from GTEx and scRNA-seq. J.W. helped with discussion and interpretation of results. H.M and J.H wrote the manuscript. All authors provided the final approval of the manuscript.

### Competing interests

**The authors declare no competing interests**.

## Supplementary figure legends

**Fig. S1.**
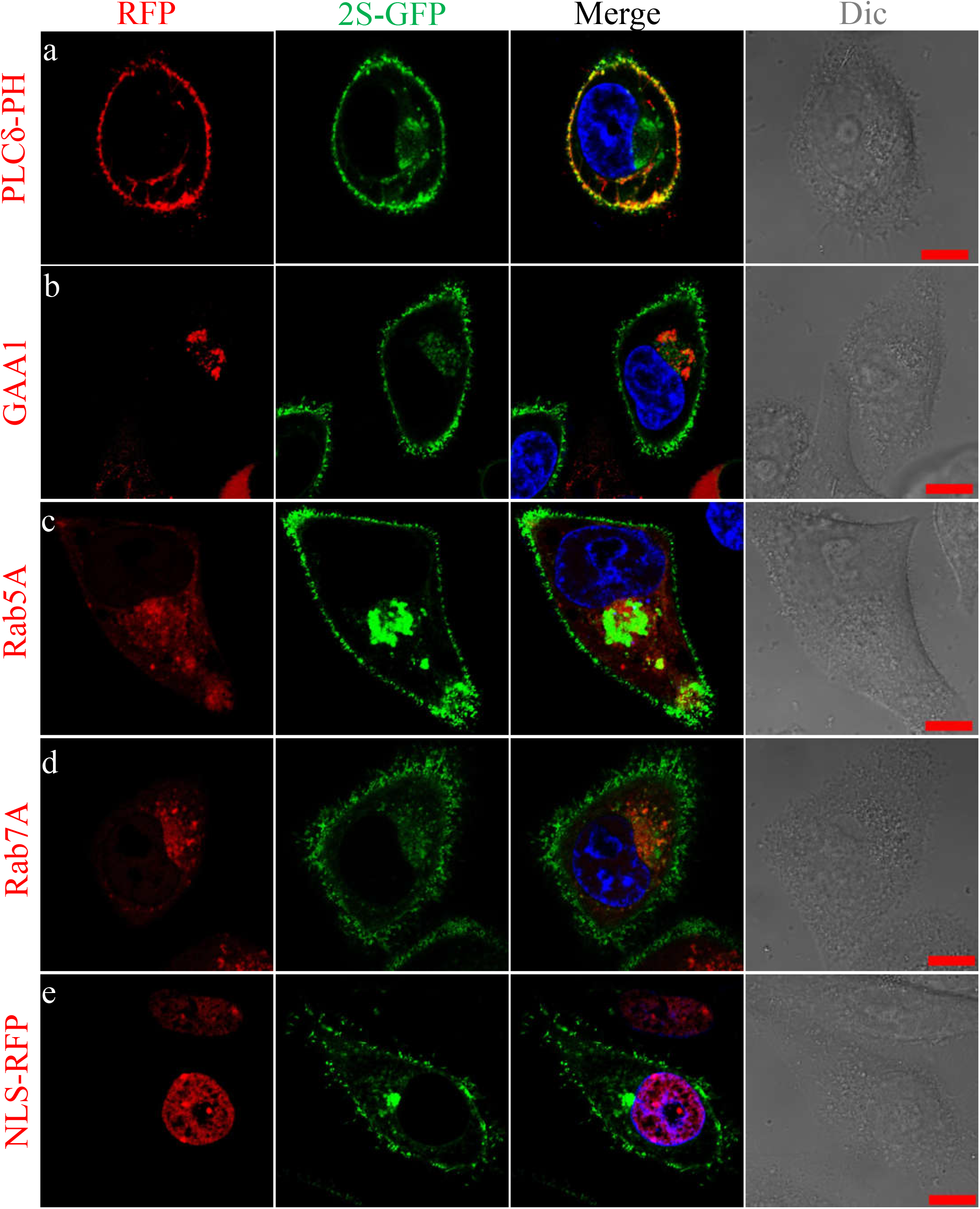
The cellular distribution of SARS-CoV-2-S-GFP. Co-expression of SARS-CoV-2-S-GFP with different markers fused with RFP in HeLa cells: (**a**). PLCδ-PH is the cell membrane marker; (**b**). GGA1 is a Golgi associated protein; (**c**). Rab5A is mainly localized in early endosome membrane or other cytosolic vesicle; (**d**). Rab7A governs early-to-late endosomal maturation; (**e**). NLS-RPF was referred as insert the nuclear localization sequence (NLS) of SV40 at the head of RPF. The images were scanned by LSM780, the nucleus(blue) was stained by Hoechst; Bar, 20 μm.

**Fig. S2:**
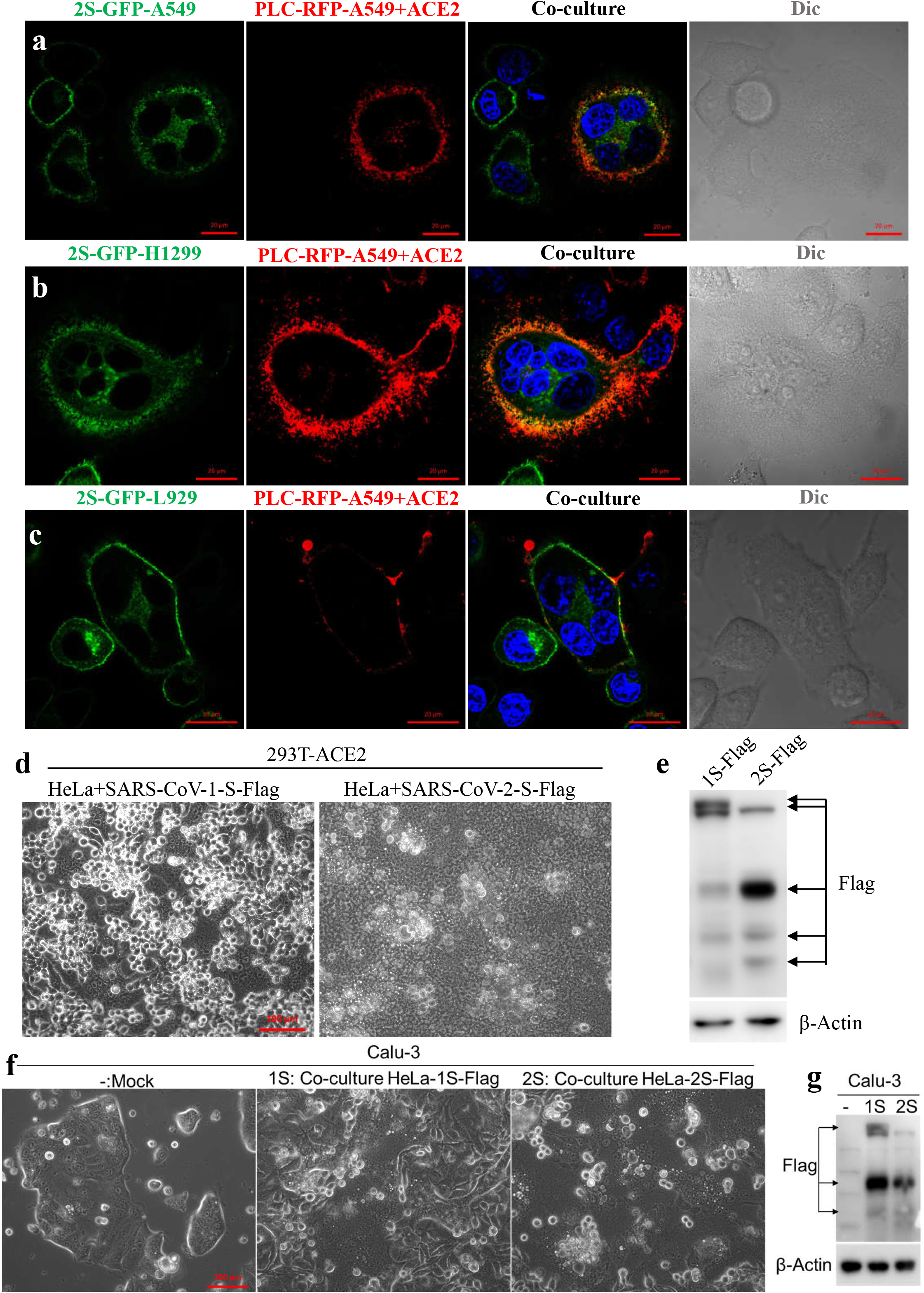
PLC-RFP-A549+ACE2 cells were co-cultured with SARS-CoV-2-S-GFP-A549 cells (**a**), SARS-CoV-2-S-GFP-H1299 cells (**b**), SARS-CoV-2-S-GFP-L929 cells (**c**) at a 1:1 ratio, four hours later, image was scanned by LSM780, the nucleus(blue) was stained by Hoechst; Bar, 20 μm. HeLa-SARS-CoV-1-S, or HeLa-SARS-CoV-2-S cells were co-cultured with 293T-ACE2 cells (**d**), or Calu-3 cells (**f**), four hours later, images were obtained by Axio Observer 5; Bar, 100 μm. Western bolt analyzed the samples collected as indicated (**e** and **g**), anti-Flag and β-Actin were probed.

**Fig. S3:**
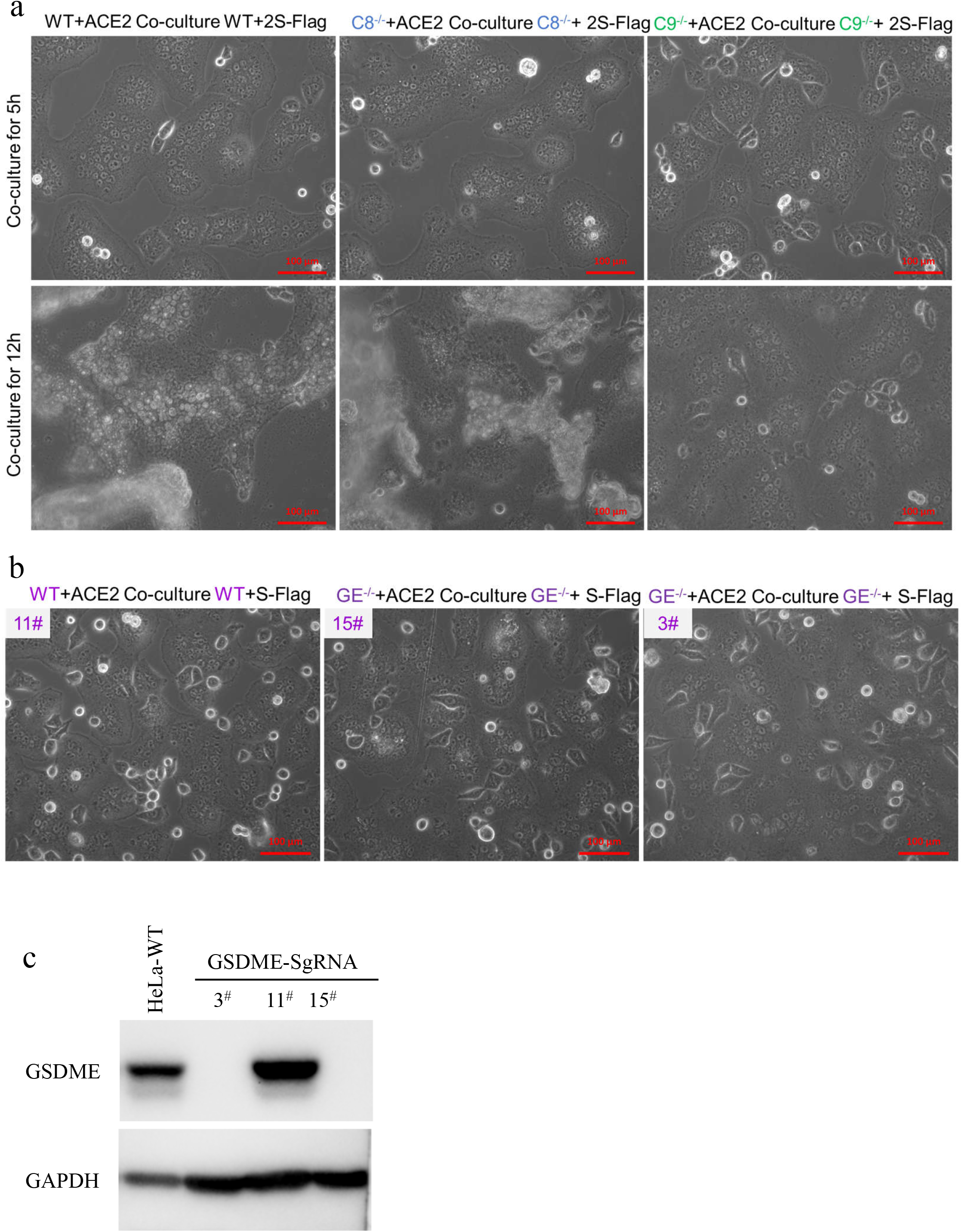
a. Generated Caspase-8(*Casp8*^*-/-*^), Caspase-9(*Casp9*^*-/-*^) knock-out HeLa cells by CRISPR-Cas9; then transfected these cells with ACE2, or SARS-CoV-2-S-Flag to obtained wild-type HeLa (WT+ACE2; WT+2S-Flag); Caspase-8 knock-out (*Casp8*^*-/-*^+ACE2; *Casp8*^*-/-*^+2S-Flag); Caspase-9 knock-out (*Casp9*^*-/-*^+ACE2; *Casp9*^*-/-*^+2S-Flag); then co-culture these cells, images were obtained by Axio Observer at 5 hours indicated for syncytia formation, and 12 hours for cell death; Bar, 100 μm. b. Images were obtained by Axio Observer at 5 hours indicated for syncytia formation as indicated; Bar, 100 μm. c. Western blot analyzed the GSDME-KO cell lines.

**Fig. S4.**
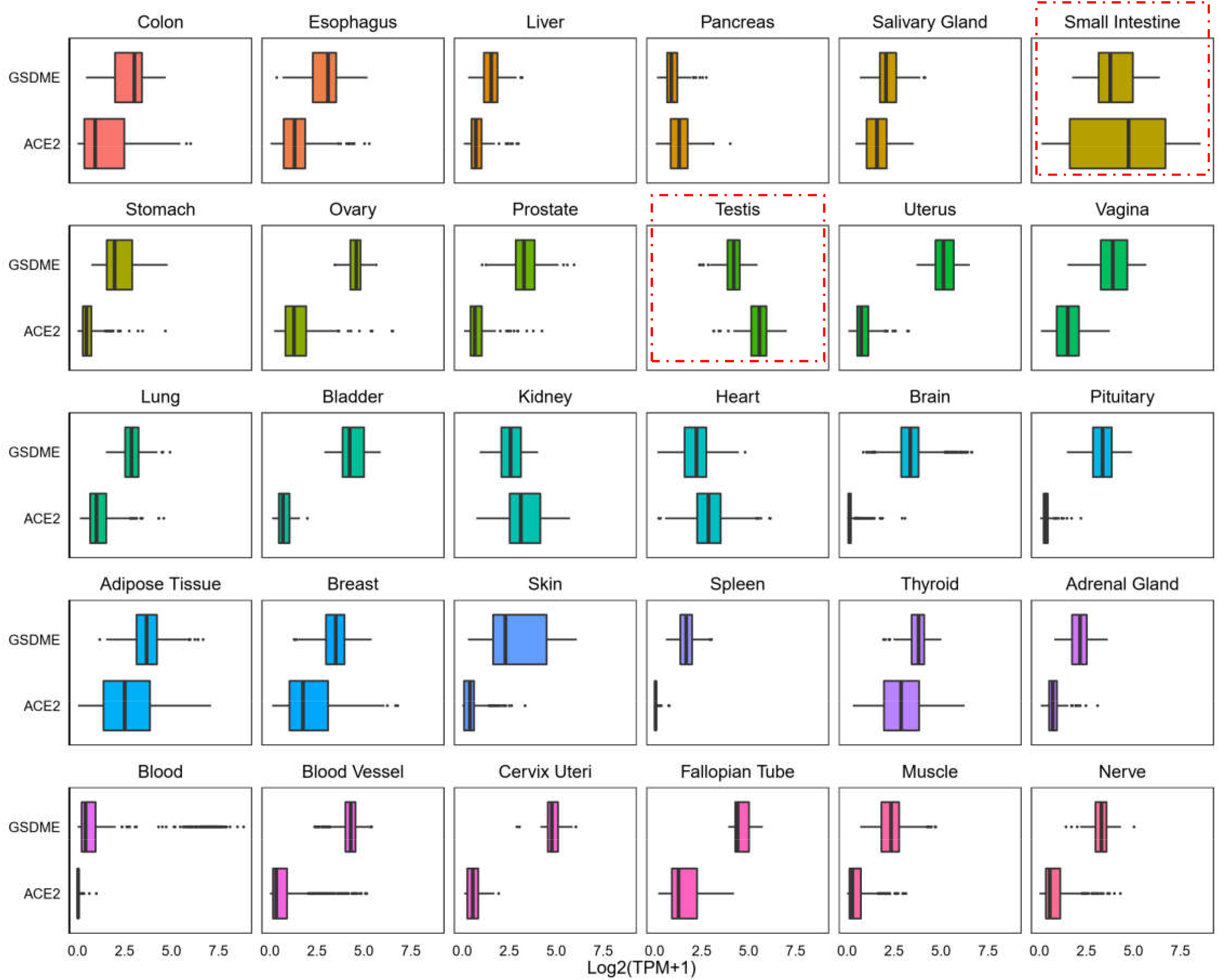
The expression profile of GSDME and ACE2. Data from the Genotype-Tissue Expression (GTEx) Project for tissues.

**Fig. S5.**
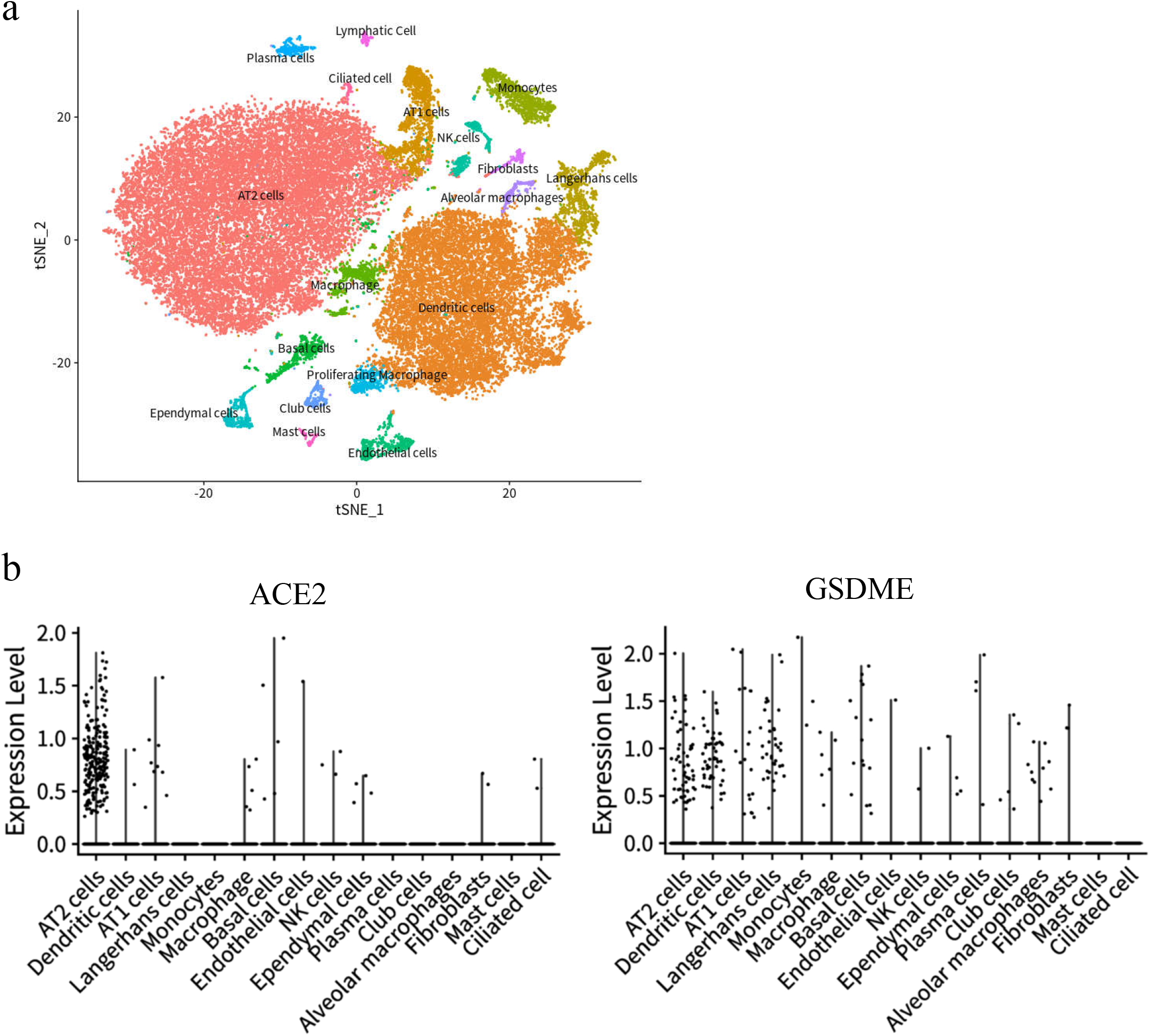
scRNA-seq profile of GSDME and ACE2 in human lung. a. Cellular populations identified. Cells were clustered using a graph-based shared nearest neighbor clustering approach and visualized using a t-distributed Stochastic Neighbor Embedding (tSNE) plot. AT1 = alveolar type □ AT2 = alveolar type □. b. Violin plots representing expression probability distributions across clusters.

## Notes

### Competing Interest Statement

The authors have declared no competing interest.

