## Supplementary material for "Pyroptosis of syncytia formed by fusion of SARS-CoV-2 Spike and ACE2 expressing cells": Methods and materials

**Plasmids and cell lines**

The plasmids of pCDNA6B-Flag-SARS-CoV-2-S and SARS-CoV-1-S provided by Prof. Wang Peihui (Shangdong University) were used as PCR template, and subcloned into pBOB-C-GFP, or pBOB-C-Flag lentivirus vector by the exonuclease III (Exo III)-assisted ligase-free cloning method; ACE2, TMPRSS2 were amplified by PCR from our human cDNA library, and subcloned into pBOB, or pBOB-C-Flag lentivirus vector; The RFP-tagged organelle markers constructs were provided by Prof. Chen Xin (Xiamen University).

HeLa, A549, H1299 and HEK293T were purchased from ATCC, and kept in Han lab at Xiamen university; Calu-3 (Procell CL-0054) were kindly provided by Procell Life Science &Technology Co., Ltd. All these cells were cultured in Dulbecco’s modified Eagle’s medium (DMEM), or Minimum Essential Medium Eagle (MEM, Calu-3) supplemented with 10% fetal bovine serum, 2 mM L-glutamine, 100 IU penicillin, and 100 mg/ml streptomycin and were kept at 37°C in a humidified atmosphere containing 5% CO2.

**Antibodies and Reagents**

The following antibodies were obtained from Proteintech: Caspase-8(13423-1-AP), Caspase-9(10380-1-AP), Caspase-3(19677-1-AP), Caspase-7(27155-1-AP), and ACE2(21115-1-AP); Cleaved-Caspase-3(9664) was purchased from Cell Signaling; DYKDDDDK-Tag(3B9) Mouse Antibody was purchased from Ab-mart; Anti-DFNA5/GSDME antibody(ab215191) was purchased from Abcam.

Chloroquine (CQ) and Camostat mesylate were obtained from MCE; The pan-caspase fmk inhibitor ZVAD was purchased from R&D Systems; Hoechst 33258 and LDH Cytotoxicity Assay Kit were purchased from beyotime; Cell Titer-Glo Luminescent Cell Viability Assay Kit was obtained from Promega.

**Confocal microscopy**

All the images were obtained in live-cell scanning model. The cells harboring GFP or RFP tagged plasmid were cultured in 35mm Glass Bottom Culture Dishes (Nest), at the indicated time. Imaging was carried out using the Zeiss LSM 780 with a 100x/1.49 NA oil objective in a 37°C incubator containing 5% CO2. GFP was excited under a 488-nm argon laser, and RFP was excited under a 568-nm argon laser. Nuclei were stained using Hoechst 33342 (1:10,000) and excited under a 405-nm argon laser.

Cell-cell fusion assay

**Western blot**

The indicated cells were harvested and immediately lysed with 1.2x SDS sample buffer. Total cell lysates were separated by SDS-PAGE and transferred to polyvinylidene fluoride membranes (EMD Millipore, 0.22μm) for Western blotting analysis with the appropriate antibodies. The proteins were visualized by enhanced chemiluminescence in accordance with the manufacturer’s instructions (NcmECL Ultra, Enhanced Chemiluminescent, ECL).

**Analysis of scRNA-seq data**

We analyzed a dataset (Gene Expression Omnibus GSE122960) freely available from public database. We used the R (version 3.6.3) and Seurat R package (version 3.9.9.9010). Firstly, Low-quality cells with less than 200 detected genes and genes found to be expressed in less than three cells were removed. Then, according to the median number of genes and the percentage of mitochondrial genes in the lung samples, we filtered cells that have unique feature counts over 5000 and cells that have >10% mitochondrial counts. After QC (Quality control), 42225 high quality lung cells were obtained.

For each individual sample, we employed a global-scaling normalization method “LogNormalize” that normalizes the feature expression measurements for each cell by the total expression, multiplies this by a scale factor (10,000 by default), and log-transforms the result. Identification of highly variable features was implemented in the FindVariableFeatures function. We detected the batch effect between eight different lung samples. So, we used the functions (FindIntegrationAnchors and IntegrateData) to integrate all eight individual samples and mitigate batch effects. Mitochondrial genes and differences in cell cycle stages between proliferating cells were regressed out by specifying the vars.to.regress argument in Seurat function ScaleData. We performed PCA on the scaled data, only the previously determined variable features were used as input. Identification of the true dimensionality of the dataset for further analysis was constructed by generating an “Elbow plot”.

To cluster the cells, we first used the FindNeighbors function to construct a KNN graph based on the euclidean distance in PCA space, and refine the edge weights between any two cells based on the shared overlap in their local neighborhoods. We next iteratively grouped cells together by using FindClusters function in Seurat. With a resolution of 0.2, cells were clustered and classified into 18 different cell types. The process to identify the cluster marker genes was implemented by the function FindAllMarkers in Seurat. Referring to three databases (CellMarker; PanglaoDB-A Single Cell Sequencing Resource For Gene Expression Data; The Human Protein Atlas), cell type assignment was performed based on the marker genes). We visualized the dataset with tSNE (t-distributed stochastic neighbor embedding) a non-linear dimensional reduction technique.
